# Dissecting the coma spectrum using Bayesian classification

**DOI:** 10.1101/832824

**Authors:** Martin J. Dietz, Bochra Zareini, Risto Näätänen, Morten Overgaard

**Affiliations:** Center of Functionally Integrative Neuroscience, Institute of Clinical Medicine, Aarhus University, Denmark; Institute of Psychology, University of Tartu, Estonia

**Keywords:** Coma spectrum, EEG, Bayesian model selection, Classification

## Abstract

A patient who does not regain full consciousness after coma is typically classified as being in a vegetative state or a minimally conscious state. While the key determinants in this differential diagnosis are inferred uniquely from the observed behaviour of the patient, nothing can, in principle, be known about the patient’s awareness of the external world. Given the subjective nature of current diagnostic practice, the quest for neurophysiological markers that could complement the nosology of the coma spectrum is becoming more and more acute. We here present a method for the classification of patients based on electrophysiological responses using Bayesian model selection. We validate the method in a sample of fourteen patients with a clinical disorder of consciousness (DoC) and a control group of fifteen healthy adults. By formally comparing a set of alternative hypotheses about the nosology of DoC patients, the results of our validation study show that we can disambiguate between alternative models of how patients are classified. Although limited to this small sample of patients, this allowed us to assert that there is no evidence of subgroups when looking at the MMN response in this sample of patients. We believe that the methods presented in this article are an important contribution to testing alternative hypotheses about how patients are grouped at both the group and single-patient level and propose that electrophysiological responses, recorded invasively or non-invasively, may be informative for the nosology of the coma spectrum on a par with behavioural diagnosis.

## Introduction

Patients with a pathological impairment of consciousness following coma are typically classified as being in either of two clinical states according to the Coma Recovery Scale (Giacino *et al.*, 2004): the vegetative state (VS) or the minimally conscious state (MCS). Currently, the two key determinants in this differential diagnosis are the patient’s sleep-wake cycle and whether motor behaviour is believed to be voluntary or simply reflexive. While the comatose patient shows no sign of arousal and is believed to have no awareness of the external world, the transition from coma to the vegetative state involves a recovery of arousal as evidenced by eye opening and a regular sleep-wake cycle. Yet, motor behaviour remains only reflexive with no signs of voluntary behaviour. Based on this observation, vegetative patients are generally believed to be as unaware of external stimuli as coma patients. Minimally conscious patients, on the other hand, are believed to show a fluctuation in their level of consciousness. They seem able to perform voluntary actions, yet often inconsistently. Nonetheless, they show no sign of verbal or behavioural communication (Laureys and Boly, 2008). It is important to emphasize that the concept of “consciousness” is used in a clinical sense to denote the assessment of a patient and not the subjective conscious experience of the patient, which in principle cannot be known. These two definitions are often confused in the literature, where the former is taken as direct evidence for the latter, although there is no scientific evidence to support such a claim (M. Overgaard, 2009; M. Overgaard and R. Overgaard, 2011). Despite the use of a standardized diagnostic scale, the subjective nature of the behavioural assessment is evident from the large variability in diagnostic consistency between health professionals, where errors are reported to range from 30-40 % and can have adverse consequences for decisions about treatment (Andrews *et al.*, 1996; Schnakers *et al.*, 2009). In the light of this diagnostic uncertainty, the desire for more objective physiological markers that can complement the nosology of the coma spectrum is becoming more and more acute. While the diagnostic determinants that define a VS or MCS patient are inferred uniquely from the patient’s behaviour, nothing can, in principle, be known about the patient’s awareness of the external world. This is because the nature of their condition precludes any report of stimulus awareness through standardised neuropsychological or psychophysical testing (M. Overgaard and R. Overgaard, 2011). This means that we have to infer a dysfunction of the neurophysiological mechanisms that mediate perception of the external world by comparison to a cohort with voluntary behaviour and normal perceptual awareness. With recent advances in neuroimaging and electrophysiological techniques, we believe that electrophysiological responses, recorded invasively or non-invasively, can serve as an informative complement to behavioural diagnostics in the classification of patients with a disorder of consciousness. In particular, the mismatch negativity (MMN), a well characterised event-related potential (ERP) to irregular changes in auditory stimulation, has been proposed as one such physiological marker (Fischer *et al.*, 1999; Näätänen *et al.*, 2011). The MMN has been used as a predictor of awakening from coma (Fischer *et al.*, 2004; 2006) and recovery from the persistent vegetative state (Wijnen *et al.*, 2007). Furthermore, the MMN is thought to be informative in predicting whether or not a comatose patient will transition to the vegetative state (Fischer and Luauté, 2005) and may even be used to differentiate the minimally conscious state from the vegetative state (van der Stelt and van Boxtel, 2008).

However, no framework has yet been proposed that enables the classification of patients at both the group level and the single-patient level. We therefore describe the use of model selection to the problem of classifying a single patient in relation to one or more nosological classes, based on their electrophysiological data. Unlike previous studies in the coma literature, this allows us to explicitly compare alternative models of patient nosology without assuming that the classification of the CRS-r is authoritative when it comes to the intactness of a patient’s sensory organs. This approach proceeds in two stages and begins with selecting an optimal model among a set of alternative hypotheses about how patients are grouped at the second (group) level. Model selection is based on the log-likelihood of each alternative model, known as the Bayesian model evidence. One could stop here and simply make an inference about how patients and controls are optimally group at the second level, under the Bayesian model evidence. However, in clinical research we are usually interested in assigning a probability of class membership to each single patient. In a second step, the set of alternative hypotheses are then used as training models, in a supervised setting, to classify each single patient and healthy control using a leave-*one*-out scheme. In this paper we used a quadratic discriminant analysis to assign a posterior probability of class membership to each patient. However, any other probabilistic model may serve, such as multinomial logistic regression (Bishop, 2006) or a neural network (Vieira *et al.*, 2017). While the group analysis is a mass-univariate analysis of the entire dataset over post-stimulus time resulting in a posterior probability map (Penny and Ridgway, 2013), the classification of each single patient (and healthy control) proceeds as a multivariate analysis, mapping from multiple data features over post-stimulus to a single probability of class membership. We validate the method in a sample of fourteen patients with a clinical disorder of consciousness and a control group of fifteen healthy adults by analysing the sensitivity and specificity of each alternative model of the nosology of patients with a disorder of consciousness.

## Materials and methods

### Patients

We recruited fourteen patients with a clinical diagnosis of coma, vegetative state or minimal conscious state following brain damage. The time since the neurological event for each of these patients had a natural progression as comatose patients will enter the vegetative state after 2-4 weeks and later progress to the minimally conscious state. To ensure the generalizability of our results, we included patients with different aetiologies and clinical histories to characterise commonalities in awareness of the sensorium. The EEG data were acquired while all patients were in an un-sedated condition following ethical approval from the local ethics committee of the Central Denmark Region, Denmark. Written informed consent was obtained from their legal surrogate.

### Coma

Five coma patients (four women) were recruited from the Department of Anaesthesiology and Intensive Care, Aarhus University Hospital, Aarhus, Denmark. Patients were included irrespective of aetiology. Written informed consent was obtained from their legal surrogate. Patients were excluded from the study if they had been declared brain dead or had a Glasgow Coma Scale (GCS) score above 9 after ending the sedative treatment. Two patients suffered from intracranial haemorrhage and three from subarachnoid bleeding. Their mean age was 71 years (range 57 to 79 years). All patients had a GCS score below 9 (range 3 to 8) after 24 hours without sedatives. The time between coma onset and EEG data acquisition was 13 days on average (range 5 to 20 days). One patient was recently diagnosed with Parkinson’s disease, but had not begun medical treatment. One patient had experienced a stroke at young age. None of the other patients had previously suffered from physical or mental illness. All patients had a Glasgow Coma score below 9 (range 3-8) after 24 hours in an un-sedated state.

### Vegetative state

Five patients (three women) in the vegetative state were recruited were recruited from Hammel Neurorehabilitation and Research Centre, Aarhus University Hospital, Denmark. The vegetative state included the following aetiologies: trauma (1), subarachnoid bleeding (2), anoxia following prolonged cardiac arrest (1) and hypoglycaemic injury (1). Their mean age was 44 years (range 30 to 65 years) and the average number days form injury to EEG acquisition was 69 days (range 46 to 131 days). These patients were examined with the Norwegian adaptation of the Coma Recovery Scale revised (CRS-r) by a trained ergotherapist and a medical student training in neurology. Most of the patients underwent two serial examinations, either on the same day or on two consecutive days. Only two patients were only examined once. Parallel with our study, the patients underwent the CRS-r performed by neuropsychologist and a nurse as a part of their rehabilitation assessment. We used these additional assessments to confirm our own assessment of each patient.

### Minimally conscious state

Four patients (two women) in the minimally conscious state were likewise from Hammel Neurorehabilitation and Research Centre, Aarhus University Hospital, Denmark. The minimally conscious state included the following aetiologies: trauma (2), subarachnoid bleeding (1) and stroke followed by an intracranial haemorrhage (1). Their mean age was 54 years (range 51 to 78 years) and the average number days from injury to EEG acquisition was 161 (range 44 to 403 days). These patients were examined with the Norwegian adaptation of the Coma Recovery Scale revised (CRS-r) by a trained ergotherapist and a medical student training in neurology. Again, the patients underwent the CRS-r performed by neuropsychologist and a nurse as a part of their rehabilitation assessment. We used these additional assessments to confirm our own assessment of each patient.

### Healthy controls

Fifteen healthy right-handed volunteers (9 women) were recruited from the Aarhus area through an online participant database supported by Aarhus University, Denmark. Their mean age was 24 years (range 19 to 35). All participants gave their informed written consent before the experiment.

### Behavioural classification

In the acute and sub-acute stages of coma, the GCS is the most widely used scale to assess the level of consciousness. It consists of a relatively brief test with three components: eye opening, verbal response and motor response. Each component has five subscales and scoring ranges from 3 to 15. A score below 9 defines a coma patient. VS and MCS patients were classified according to the CRS-r. This scale was originally designed in 1991 at the JFK Johnson Rehabilitation Institute and later revised in 2004 by Giacino and colleagues. The purpose of this scale is to assist with differential diagnosis, prognostic assessment and treatment planning. The scale consists of six categories addressing: auditory, visual, motor and oro-motor function, communication and arousal. The subscales are comprised of a hierarchy of items, the lowest reflecting reflex activity and the highest representing cognitively mediated behaviours (Giacino *et al.*, 2004). Table 1 shows demographic details and behavioural scores. The average GCS score in the coma group was 6 (range 4 to 8). The average CRS-r score in the VS group was 5.8 (range 4 to 9). The average CRS-r score in the MCS group was 13.8 (range 11 to 20).

**Table 1.** Clinical scores indicated by the GCS and the CRS-r. SAH = Subarachnoid haemorrhage, ICH = intracerebral haemorrhage, M = male, F = female, VS = vegetative state, MCS = minimally conscious state, SD = standard deviation.

| Patient | Aetiology | Sex | Age<br>(years) | Time of EEG after<br>insult (days) | GCS | CRS-r | Nosology |
| --- | --- | --- | --- | --- | --- | --- | --- |
| 1 | Apoplexy,<br>ICH | F | 79 | 5 | 4 | - | Coma |
| 2 | ICH | F | 73 | 10 | 3 | - | Coma |
| 3 | SAH | F | 69 | 20 | 7 | - | Coma |
| 4 | SAH | M | 77 | 20 | 8 | - | Coma |
| 5 | SAH | F | 57 | 12 | 8 | - | Coma |
| Mean estimate (SD) |  |  | 71 (5.2) | 14.4 (39.6) | 6 (2.3) |  |  |
| 6 | Trauma | M | 30 | 131 | - | 9 | VS |
| 7 | SAH | F | 40 | 51 | - | 3 | VS |
| 8 | Anoxia | F | 65 | 54 | - | 7 | VS |
| 9 | SAH | M | 46 | 64 | - | 4 | VS |
| 10 | Anoxia | F | 40 | 46 | - | 6 | VS |
| Mean estimate (SD) |  |  | 44 (7.3) | 42.2 (56.0) |  | 5.8 (2.4) |  |
| 11 | Trauma | M | 51 | 124 | - | 20 | MCS |
| 12 | Apoplexy, ICH | F | 78 | 75 | - | 13 | MCS |
| 13 | Trauma | M | 53 | 403 | - | 11 | MCS |
| 14 | SAH | F | 53 | 44 | - | 11 | MCS |
| Mean estimate (SD) |  |  | 58 (7.8) | 137.7 (59.4) | - | 13.8 (4.3) |  |

### Experimental paradigm

The stimulus paradigm consisted of a sequence standard tones, each followed by a deviant tone that differed along a different auditory dimension (Näätänen et al., 2004). The stimuli were composed of harmonic tones consisting of 3 sinusoidal partials of 500, 1000 and 1500 Hz with 75 ms duration, including 5 ms fade-in and fade-out. The oddball or deviant tones differed from the standard tone in pitch, intensity, duration, location and by having a gap in the middle of the tone. Half of the pitch deviants were 10 % lower than the standard (450, 900 and 1350 Hz), while the other half were 10 % higher (550, 1100 and 1650 Hz). Half the intensity deviants were 10 dB lower, while the other half were 10 dB higher. The gap tone was created by inserting 7 ms of silence in the middle of a standard tone. The change in location was created by introducing an interaural time difference of 800 μs between the left and right ears. This corresponds to a perceived deviation in sound location of 90 degrees angle to the left or right of the subject relative to midline (Paavilainen et al., 1989). The duration deviant was 50 ms shorter than the standard tone, resulting in 25 ms duration. The deviants were delivered in semi-randomized order, so that the same type of deviant was never repeated. The stimulus-onset asynchrony was 500 ms and the total duration of the experiment was 16 min, starting with 15 standard tones. This resulted in a total of 1920 stimuli of which half were standard tones and half were deviant tones. The stimuli were presented binaurally via headphones (Sennheiser, GmbH) using Presentation software (Neurobehavioural Systems, Inc).

### Data acquisition

We recorded 32-channel electroencephalography (EEG) using active Ag/AgCl electrodes placed according to the extended 10-20% system, sampled at 1 kHz (Brain Products, GmbH). Given that the coma group could not fit the usual caps, we used a custom cap with 25 channels that conformed to the extended 10-20% system to ensure commensurability across groups. The EEG was referenced online to FCz and the electro-oculogram (EOG) was recorded from two electrodes placed above and on the outer canthus of the right eye.

### EEG data analysis

Data analysis was performed using Statistical Parametric Mapping (SPM12, revision 6685) academic software implemented in Matlab (MathWorks Inc., USA). EEG data were high-pass filtered at 0.5 Hz and low-pass filtered at 30 Hz using a two-pass Butterworth filter and down-sampled from 1 kHz to 250 Hz. Experimental trials were epoched from −100 to 400 ms in peri-stimulus time and baseline-corrected using the average over the pre-stimulus time-window. Artefacts were rejected by thresholding the signal at 80 *µ*V and re-referenced to the average over all channels. Single trials were averaged using robust averaging to form event-related responses (ERPs) that are phase-locked to the stimulus. To facilitate analysis in sensor space, the ERPs were converted into 3D spatiotemporal images (2D scalp over post-stimulus time). These images of evoked responses in each patient and control were then used as summary statistics for the Bayesian model comparison described below using Bayesian inference at the second level (Penny and Ridgway, 2013).

### Bayesian model selection for patient grouping

Model selection is based on the evidence or marginal probability of a model, in relation to a set of alternative models, given the same data. The model evidence is then used for Bayesian inference on models that encode difference hypotheses in terms of a posterior probability map (PPM) over the spatiotemporal image of electrophysiological responses. This leads to a natural interpretation of patient grouping in data space. Here, we use this approach to the problem of classifying *N* patients and healthy controls into an optimal set of distinct classes *𝒞*_*k*_, for *k = 1*,…,*K*, given the data features of their average electrophysiological responses. As we will see, optimal means the model with the highest evidence. We begin by formulating a set of models *ℳ* ∋ {*m*_1_,…, *m*_*k*_} where each model *m*_*k*_ represents an alternative hypothesis about the grouping of patients into *k* classes. These models can be formulated as a Bayesian general linear model of the form *y* = *Xϑ* + *ϵ* where data features *y* ∈ ℝ^*n*×1^ are given by the average electrophysiological response of each patient (and control) at each pixel and time bin of a 3-dimensional spatio-temporal image and *ϵ* ∈ ℝ^*n*×1^ is zero-mean additive observation error *p*(*ϵ*) = *𝒩*(*ϵ* | 0,*C*_*ϵ*_). Here, each model is given by a design matrix *X* ∈ ℝ^*n*×*k*^ of predictor variables encoding the grouping of *n* patients and controls into *k* distinct classes and *ϑ* are the model parameters that encode the between-group means and precisions that separate the classes. As every class is defined by having a different group mean and precision, *X* = *I*_*k*_ ⊗ **1**, where ⊗ is the Kronecker product of the *k x k* identity matrix and a unit vector of ones encoding the number patients or controls in the *k*th class. Fig. 1 shows the grouping of patients and controls according to three alternative hypotheses, each encoded by a different design matrix *X*. The first model represents the hypothesis that there is no difference between patients and healthy subjects. The second model represents the hypothesis that all patients have similar responses and that they are distinct from the responses in healthy controls. Finally, the third model represents the hypothesis that all patients have different responses.

**Figure 1.**
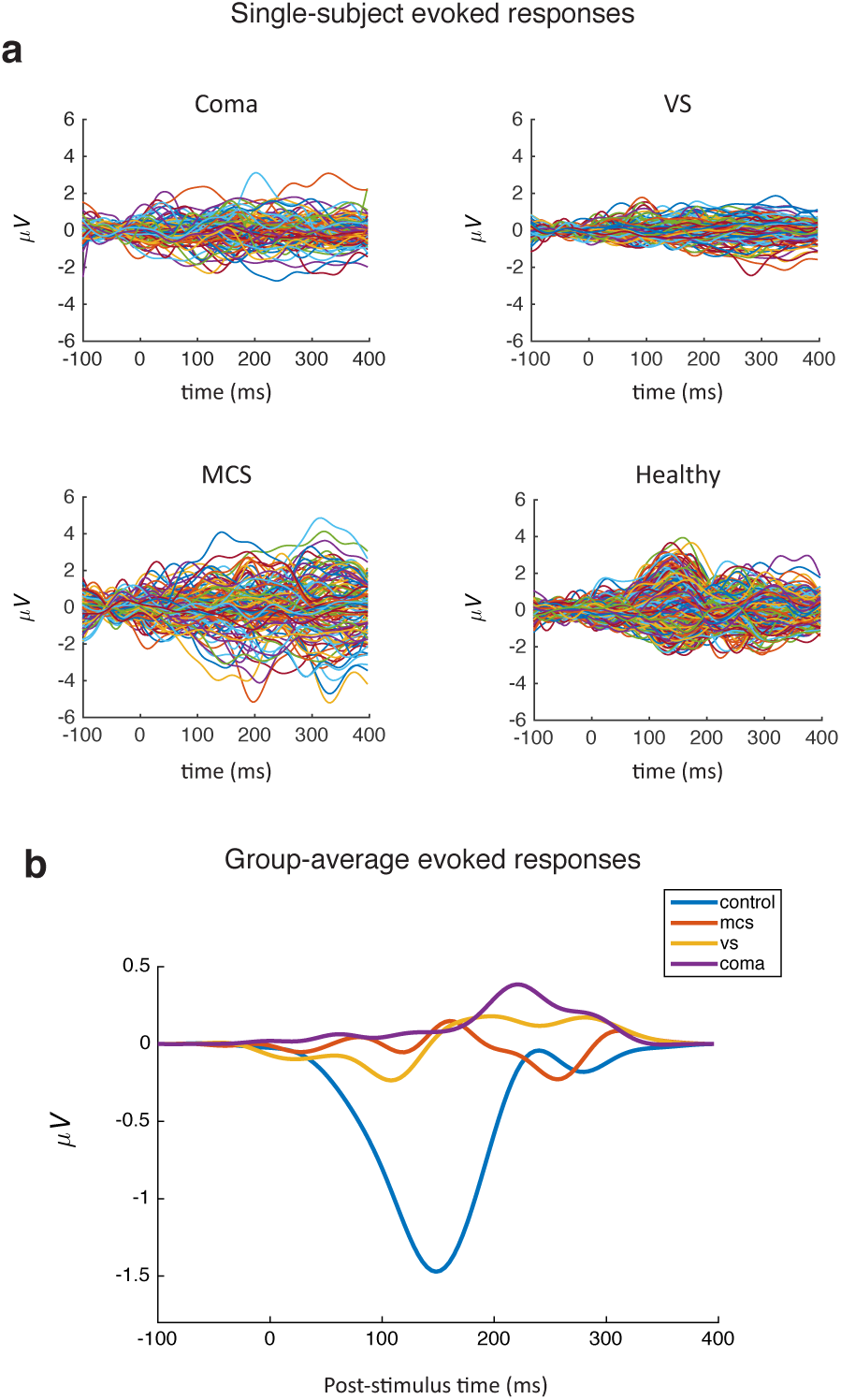
**(a)** Evoked responses to surprising stimuli, relative to repeated stimuli, at all channels over post-stimulus time. **(b)** Grand-average responses showing a classical MMN in healthy subjects and reduced responses in the patient subgroups.

For Bayesian inversion of each model, we have a multivariate Gaussian prior distribution over the model parameters at each voxel *p*(*ϑ* | *m*) = 𝒩(*ϑ* | *η*, ∑) where the prior mean is set to zero *η* = 0 and the prior covariance ∑ = *α*_*k*_^−1^*I* is estimated from all voxels in the spatiotemporal search region using empirical Bayes (Friston and Penny, 2003). This global shrinkage prior effectively shrinks the parameters towards zero in the absence of precise information in the data to inform the model. This corresponds to quadratic regularisation in classical statistics and machine learning (Bishop, 2006). The assumption behind this shrinkage prior is that, on average over all channels, the post-synaptic activity is zero, with non-zero activity expressed regionally over the scalp and regionally in post-stimulus time. This is a reasonable assumption as the data are referenced to the average over all channels and baseline-corrected to the pre-stimulus time window. The likelihood function for the data is also Gaussian and given by *p*(*y*|*ϑ, m*) = 𝒩(*y*|*Xϑ,C*_*ϵ*_) where the observation error covariance *C*_*ϵ*_ (*i*) = *λ*_*i*_*V* at the *i*th voxel is estimated from the data in terms of a voxel-specific hyperparameter *λ*_*i*_ (Penny and Ridgway, 2013). The prior and the likelihood now fully specify each model and Bayesian inversion provides the multivariate posterior distribution over the parameters of each model at each voxel *p*(*ϑ* | *y, m*) = 𝒩(*ϑ*|*μ*_*ϑ*_, *C*_*ϑ*_) with mean *μ*_*ϑ*_ and covariance *C*_*ϑ*_ given by

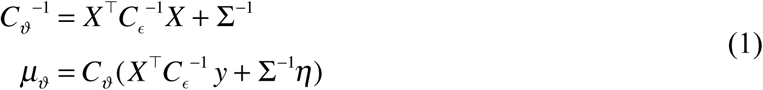

Bayesian inversion also provides and the marginal likelihood of each model itself and obtains by marginalising over the entire set of model parameters. This is known as the Bayesian model evidence

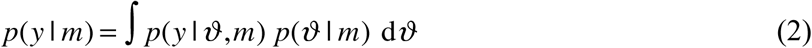

For linear models, the log-evidence ln *p*(*y* | *m*) is given by the Laplace free energy

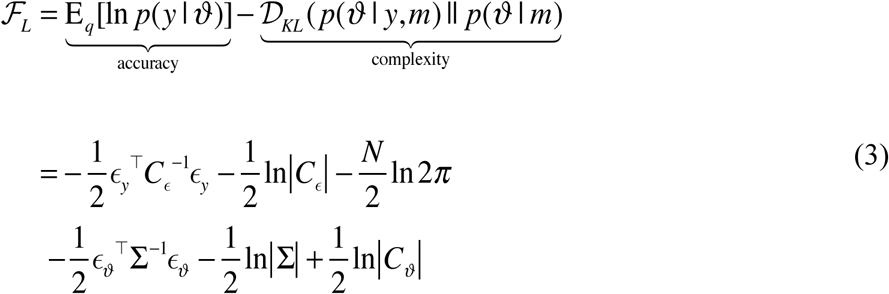

where *ϵ*_*y*_ = *y* − *Xϑ* are the prediction errors of the data, *ϵ*_*ϑ*_ = *μ*_*ϑ*_ − *η* are the parameter prediction errors and ln|·| denotes the log-determinant. We have expressed the free energy as accuracy minus complexity. Here, the accuracy is given by the expected log-likelihood of the data, given the parameters, and describes the model fit. The complexity is given by the Kullback-Leibner divergence of the posterior density from the prior density. This shows how the Bayesian model evidence accounts for model complexity by penalising models whose posterior diverges from prior beliefs after observing the data.

Under the Neyman-Pearson lemma, the most sensitive hypothesis test is the likelihood-ratio test comparing two alternative models of the data (Neyman and Pearson, 1933). In Bayesian inference, this is known as the Bayes factor (Kass and Raftery, 1995)

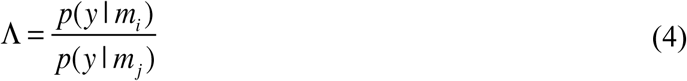

comparing the evidence of model *i* and model *j*. Unlike classical (frequentist) model scoring, the Bayesian model evidence takes into account model complexity. We used a computationally efficient approximation to the Bayes factor known as the Savage-Dickey density ratio (Dickey, 1971)

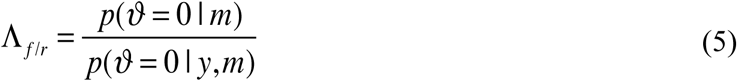

This is the ratio of the prior probability density to the posterior probability density evaluated under the null hypothesis that the parameters are exactly zero. Intuitively, this compares a full model *f* to a reduced model *r* whose parameters are exactly zero. When the prior means are *η* = 0, as is the case for the shrinkage priors used here, the log-evidence of a full model relative to a null model is given by (Penny and Ridgway, 2013) as implemented in the SPM software:

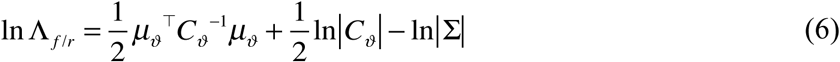

For two alternative models, their posterior probability is then given by *σ* (ln Λ), where *σ* (·) is the logistic sigmoid function

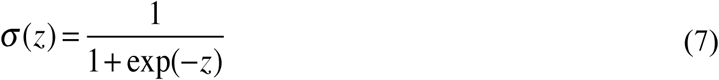

whereas the posterior probability of multiple models is given by the normalized exponential or softmax function

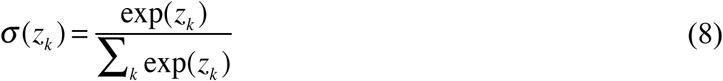

Where *z*_*k*_ = ln *p*(*y* | *m*_*k*_).

### Multivariate discriminant analysis for patient classification

For single patient classification, we used quadratic discriminant analysis in a multivariate setting. This simply requires an estimate of the mean and covariance of each training class *𝒞* _*k*_, for *k =1*…*K*, which follows from Gaussian assumptions about the observation error. However, in neuroimaging and electrophysiology we usually have many more voxels or temporal dimensions than samples, which can render the estimated covariance matrix rank-deficient. We therefore use a re-parameterization of the data covariance in terms of a set of hyperparameters that are estimated using restricted maximum likelihood (ReML) (see (Friston *et al.*, 2002) and (Friston and Penny, 2003) for an in-depth treatment in neuroimaging). The ReML estimate of the error covariance is given by *C*_*ϵ*_ = ∑*λ*_*i*_*V*_*i*_ where each hyperparameter *λ*_*i*_ is estimated for a corresponding covariance basis *V*_*i*_ that captures a particular covariance structure. In this application, we assumed an IID temporal covariance over post-stimulus time *C*_*k*_ = *λ*_*k*_*I* ^*mxm*^. Although this model assumes temporal stationarity, one could also use multiple hyperparameters over post-stimulus time to accommodate non-stationarity. Once we have estimated the means *μ*_*k*_ and covariances *C*_*k*_ of the m-dimensional feature space for each class, the discriminant functions are given by

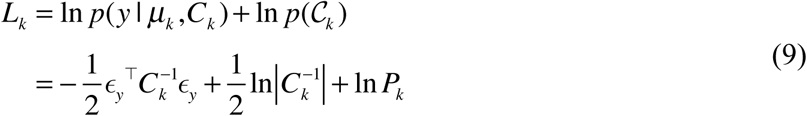

where *y* is the m-dimensional vector of data features from the *i*th subject, *ϵ*_*y*_ = *y* − *μ*_*k*_ are the prediction errors and *P*_*k*_ is the prior probability that subject *i* is a member of class *k*. The decision functions that specify the boundaries between classes are then given by the differences of discriminant functions and the posterior probability that the *i*th subject is sampled from class *k* is given in the usual way by the sigmoid or softmax function *p*(𝒞_*k*_ | *y*) = *σ* (*L*_*k*_) using equation 7 or 8. We see that the discriminant function for each class is composed of a Gaussian log-likelihood function plus a log-prior, hence the quadratic form of the decision functions. The posterior probability is then effectively given by comparing the data under different class-specific training models and the classification can therefore be viewed as a simple case of Bayesian model selection. In this application, we considered two alternative training models that each embodied an alternative hypothesis about the number of patient subgroups. The first model had two classes corresponding to patients and healthy controls, whereas the second model had four classes corresponding to three patient subgroups and healthy controls. For each alternative model of DoC nosology, we used a leave-*one*-out scheme for assigning a posterior probability that each patient (and control) comes from a given nosological class. This provides classical model diagnostics of sensitivity and specificity in terms of the classification accuracy and the receiver operating characteristic (ROC) curve. Our code for multivariate classification is available at https://github.com/martinjdietz/Bayesian-classification.

## Results

### Evoked cortical responses

We used the average electrophysiological responses from each patient and control as data features. The evoked responses from each patient group, classified according to the Coma Recovery Scale revisited (Giacino *et al.*, 2004), are shown in **Fig. 1a** and the group average in shown in **Fig. 1b** at the scalp location that showed the maximal effect of surprising stimuli, relative to repeated stimuli, over all patients and controls at 160 ms post-stimulus time, *T*_28_ = 4.59, *p* = 0.003, family-wise error corrected using Random field theory (Kilner and Friston, 2010). This showed the presence of a classical mismatch negativity (MMN) response is the dataset, without making any assumptions about differences between patients (and controls) before proceeding to the formal test of alternative hypotheses about patient classification.

### Optimal patient grouping

We then compared our alternative hypotheses about how patients are grouped in terms of their posterior probability map over the scalp and over post-stimulus time. This showed that the optimal grouping of patients was given by the model that groups comatose patients, patients in the vegetative state (VS) and patients in the minimally conscious state (MCS) as a joint class, given their electrophysiological responses (**Fig. 2a**). The posterior probability map (PPM) showed that the effect was expressed over the central scalp with a peak at 152 ms (*P* > 0.99) (**Fig. 2b**). Finally, we used a contrast of the posterior parameters of the model to obtain the posterior probability that the groups were different from each other in terms of their average responses at 152 ms post-stimulus time (**Fig. 2c**). This revealed that there is a difference between patients and healthy subjects with a high degree of confidence (*P* > 0.99).

**Figure 2.**
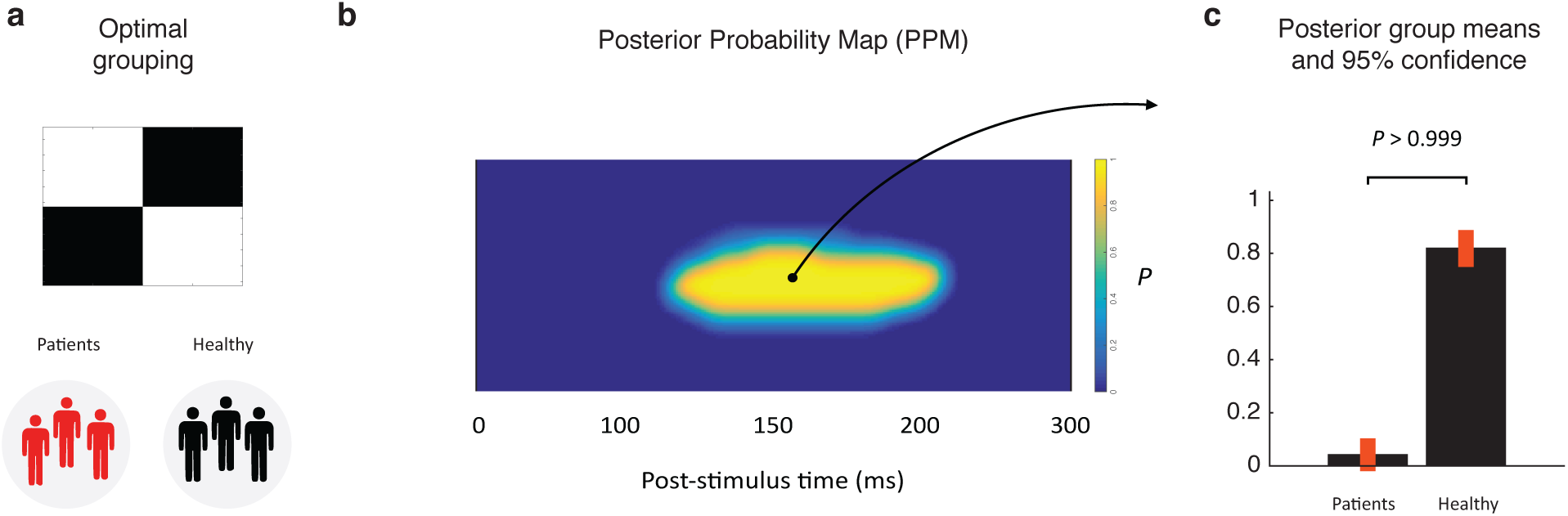
Optimal model among the set of alternative hypotheses about patient grouping **(a)** Design matrix of the optimal model encoding the partition of patients and controls into distinct classes **(b)** Posterior probability map (PPM) of the optimal model showing the difference between the two groups at 152 ms post-stimulus time, the latency of the classical mismatch negativity **(c)** Posterior group means and 95% confidence intervals.

Finally, to see why the model based on the CRS-r diagnosis of patients had lower evidence than the model treating VS and MCS patients as a joint class, we inspected the posterior parameter estimates of the VS and the MCS patients when modelled as separate groups using the third model in Fig. 1. This showed that the parameters encoding the group mean and precision of the VS and MCS patients are statistically indistinguishable from one another. Their absolute effect sizes are virtually identical and the posterior probability that they are different is *P* = 0.51 (Fig. 3b). In other words, there is no information in the MMN response to inform a difference between the two patient groups. We the inspected the correlation between the behavioural CRS-r scores, used to clinically differentiate between VS and MCS patients, and the MMN response at 152 ms. Again, there is virtually no relation between the CRS-r diagnostic scale and our neurophysiological proxy for perceptual awareness (*r* = 0.3, *p* = 0.32, linear correlation) (Fig. 3c).

**Figure 3.**
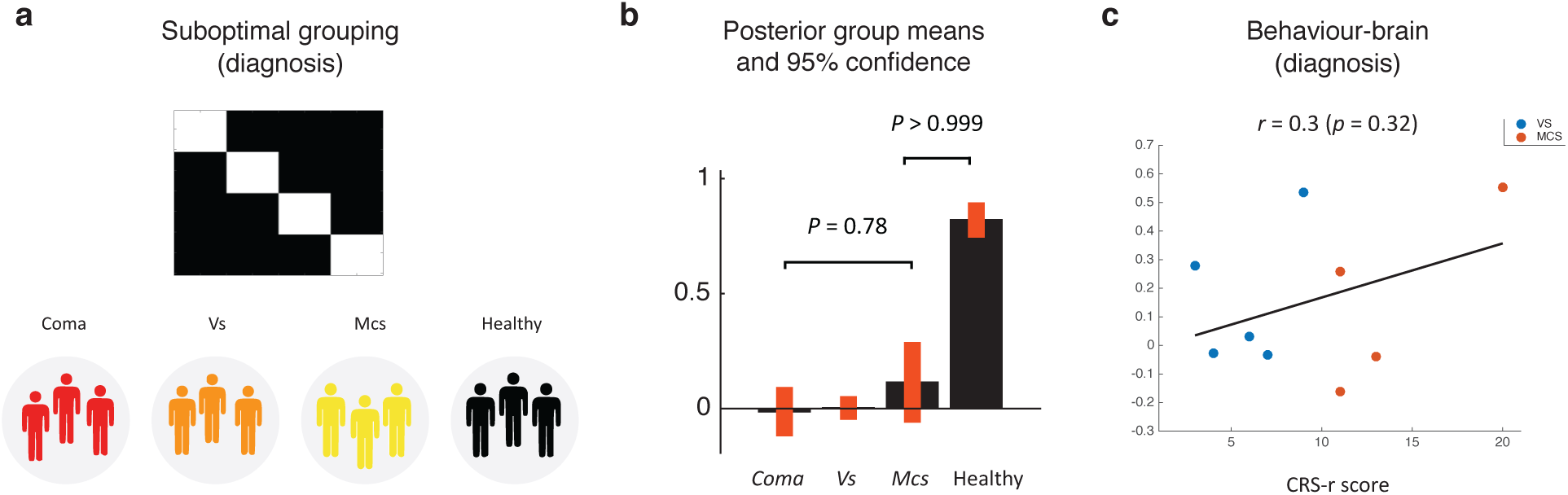
Suboptimal model of electrophysiological responses **(a)** Design matrix encoding a partition between coma, VS and MCS patients **(b)** Posterior group means and 95% confidence intervals indicating that coma, VS and MCS patients are indistinguishable at the group level **(c)** Negligible relation between CRS-r scores in VS and MCS patients and their MMN response.

### Single patient classification

Using a leave-one-out scheme for classification, we then considered two alternative training models that each embodied an alternative hypothesis about the number of patient subgroups. The first model had two classes corresponding to patients and healthy controls, whereas the second model had four classes corresponding to the three patient subgroups assumed by the Coma Recovery Scale revisited: coma, vegetative state and minimally conscious state. The data features that entered these models were selected using an *F*-test of the overall oddball response to surprising stimuli inclusing all patients and controls, thresholded at *p* < 0.05, corrected for multiple comparisons using random field theory. This resulted in a mask over channel C4 between 100 – 200 ms post-stimulus time. This approach to feature selection is known as a “collapsed localiser” (Luck and Gaspelin, 2017) and represents an unbiased way of identifying informative data features, because the *F*-test of the overall average response is orthogonal to any differences between groups (Christensen, 2011). For each of the two alternative nosological models, we used a leave-*one*-out scheme for assigning a posterior probability that each patient (and control) comes from a given class. In this application, a posterior probability *P* > 0.99 was used as the criterion for a correct classification. The classification accuracy of the model with one joint patient group was 90 %, whereas the accuracy of the model with four patient subgroups, as defined by their CRS-r scores, was 48 %. This shows that the model consisting of one joint patient group was much more accurate in predicting each patient and healthy control, resulting in the higher sensitivity to patients and the higher specificity in discerning patients from healthy controls, compared to the model of three subgroups as defined by the CRS-r. This relationship between the true positive rate (sensitivity) and the false positive rate (1 – specificity) is depicted in Fig. 4 as the receiver operating characteristic (ROC) curves for the two alternative models under a varying decision function (posterior probability criterion).

**Figure 4.**
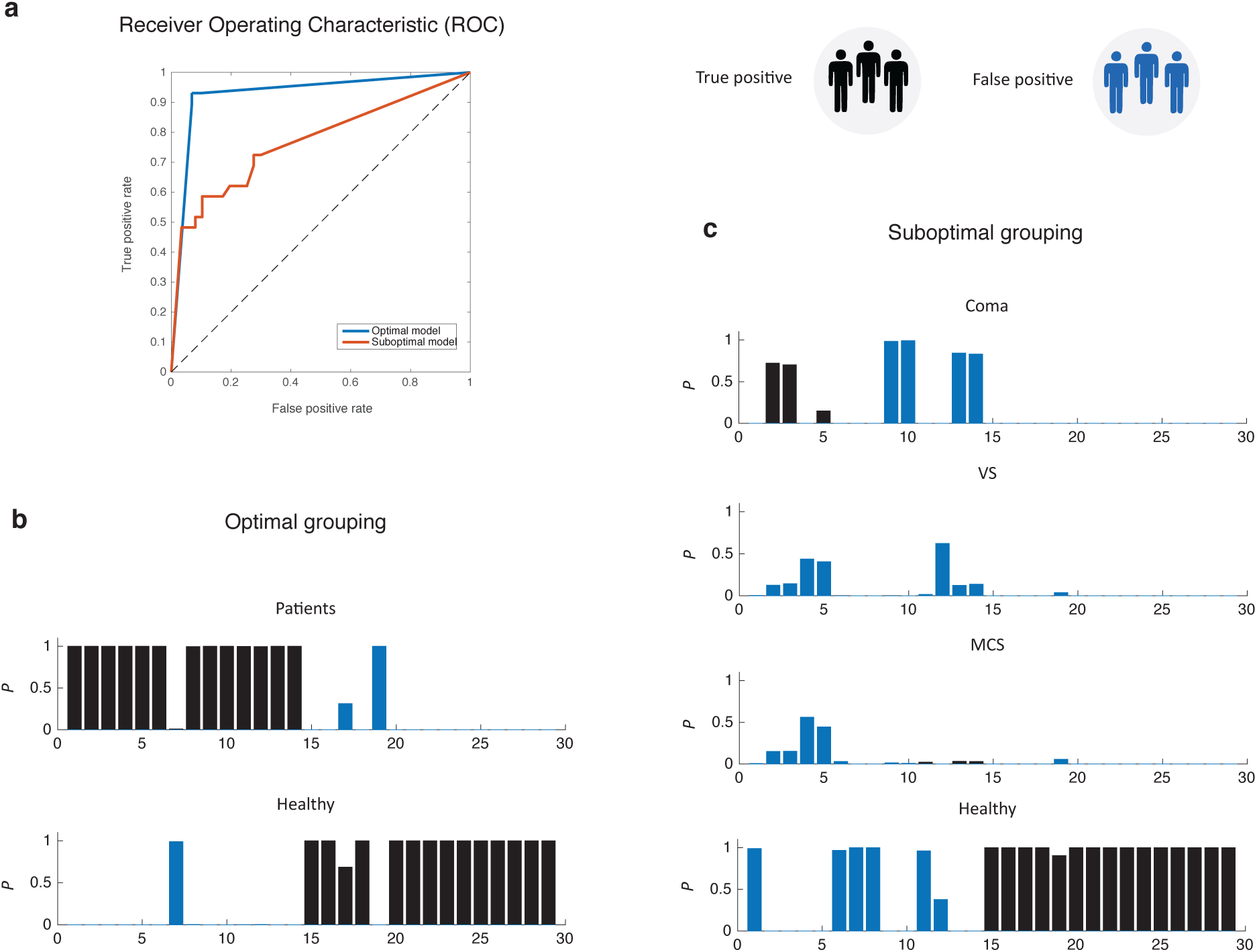
Model diagnostics of sensitivity and specificity **(a)** Receiver operating characteristic (ROC) curve of the two alternative models of patient classification **(b)** Posterior probability of each subject belonging to either the patient class or to the healthy controls **(c)** Posterior probability of each subject belonging to either of the three subclasses of the CRS-r or to the healthy controls. The probability mass functions in black are true positives and those in blue are false positives, under each alternative model of patient classification.

## Discussion

Motivated by the quest for more objective neurophysiological markers within the coma spectrum, we have described the use of Bayesian model selection to the problem of adjudicating between alternative models of how patients are grouped, given their electrophysiological data. This level of inference allows us to map between a particular patient grouping and their corresponding behavioural scores (CRS-r scores) to inform the nosology of the coma spectrum in terms of the patients’ average responses. However, this does not address the problem of classifying a single patient into one or more nosological classes. This is done using the kind of probabilistic generative model presented here, which allows a mapping from multiple data features over post-stimulus to a single probability of class membership. This approach is clearly motivated by the desire for more objective physiological markers that can complement the existing behavioural diagnostics. We explicitly compare alternative models of patient grouping without assuming that the classification of the CRS-r is authoritative when it comes to a patient’s sensory perception. This is important because the current diagnostic determinants are inferred uniquely from the observed behaviour of the patient, where nothing can be known about the patient’s awareness of the external world through his or her sensory organs. The results of our validation study show that we can disambiguate between alternative models of how patients are classified. Although limited to this small sample of patients, this allowed us to assert that there is no evidence of subgroups when looking at the MMN response in this sample of patients.

### The effects of neuropathology on electrophysiology

We used an auditory oddball paradigm as the mismatch negativity (MMN) has been proposed in the literature as a neurophysiological marker for cognitive function in patients with a disorder of consciousness. Given the probabilistic nature of oddball paradigms, they are known to induce short-term cortical plasticity at different levels of the sensory system, depending on stimulus complexity. Unexpected or surprising stimuli have consistently been shown to induce a change in cortical connectivity between lower-level sensory areas and higher-level frontal and parietal areas, both in the auditory system (Auksztulewicz and Friston, 2015; Dietz *et al.*, 2014; Garrido *et al.*, 2009) and the tactile system (Fardo *et al.*, 2017). This short-term plasticity is thought to mediate perceptual learning of sensory stimuli and is observed in the healthy brain as the MMN response. Using the same kind of oddball paradigm and dynamic causal modelling, Boly and colleagues showed that feedback connectivity from the frontal to the temporal lobe was impaired in the vegetative state compared to healthy controls (Boly *et al.*, 2011). This shows that with high-quality electrophysiological data, it is possible to use the sort of probabilistic model presented here to make predictions about new patients, based on either observed responses at the data level or the parameters from a biophysical model of effective connectivity between brain regions (Brodersen *et al.*, 2014). However, one major challenge in the source modelling of electromagnetic responses from brain-injured patients is that they have tissue damage, tissue swelling and intra-cerebral fluid changes. These pathological alterations to the neuronal tissue are likely to change their functional anatomy in an idiosyncratic way. Furthermore, skull fractures or openings in the skull due to craniotomy will invalidate standard forward models for EEG source reconstruction (Wolters *et al.*, 2006). This heterogeneity in functional anatomy caused by different pathologies speaks directly to the use of the multivariate analysis of electrophysiological responses. This is in contrast to the use of univariate analysis of each channel and time bin over post-stimulus time. This is because univariate analysis of electrophysiological data assumes a certain degree homogeneity in the orientation of the cortical columns that the generate a particular pattern of observed electromagnetic responses as measured with EEG and MEG (Buzsáki *et al.*, 2012). While this homogeneity of functional anatomy can enter an assumption in the statistical analysis of electromagnetic responses at the group level, it is an assumption that is most likely invalidated in a group analysis of brain-injured patients. This is why we use a multivariate scheme, as it allows for the expression of effects at different locations on the scalp EEG, different latencies over post-stimulus time or at different frequencies in time-frequency space between different patient classes.

### Future directions in single patient prediction

In this application, we have used the MMN as a data feature for single patient classification. However, we wish to stress that the choice of the MMN response as the central measure does not necessarily reflect the belief that it is a more informative proxy for a patient’s general pathophysiology than any other neuroimaging or electrophysiological data feature. Any electromagnetic response that is robustly expressed at the single-subject level could in principle enter as a data feature, under the assumption that it represents some sensorimotor or cognitive mechanism. While visual evoked responses are most practical in awake behaving subjects with no oculo-motor or visual pathologies, auditory and sensory evoked responses are by far the most practical to implement in a clinical setting. In the present context, the use of the MMN was a practical choice as we wanted to build on existing evidence (Fischer et al., 2006; 2004; Fischer and Luauté, 2005; Johnsen et al., 2017; Naccache et al., 2005; Tzovara et al., 2013; van der Stelt and van Boxtel, 2008; Wijnen et al., 2007). Furthermore, we wish to underline that regardless of which measure we use - neuronal or behavioural - we do not believe that we can make any conclusions about consciousness – i.e. the subjective experience – of patients based on these methods. For that reason, the very term “disorder of consciousness” should possibly be replaced with another that simply refers to the neural state of a patient. Our plans for future studies is thus to continue developing tools to predict the outcome of a new coma patient whose prognosis is unknown. This requires a large sample of patients whose follow-up status is known. This approach has a dual purpose: on the one hand, the data from this patient database and the accompanying database of healthy age-matched controls can be used, in combination with behavioural scores, to predict the outcome of new patients whose prognosis is unknown. At the same time, the sensitivity and specificity of this sort of predictive model will be gradually informed by the follow-up status new patients and this will in turn inform the nosology of the coma spectrum in a systematic and evidence-based way. Crucially, standardization of stimulus protocols, classification models and data sharing will be absolutely key in this future multi-site or global endeavour to efficiently predict final outcome of new patients in a coma.

## Author contributions

M.J.D., B.Z., R.N. and M.O. conceived the idea for the study. B.Z identified and characterised the patients. B.Z. acquired the EEG data in patients and healthy controls. M.J.D analysed the data and contributed the methodological tools. M.J.D, B.Z. and M.O. wrote the paper.

## Acknowledgments

We would like to thank Birger Johnsen and Carsten Kock-Jensen at Aarhus University Hospital, as well as Jørgen Feldbæk Nielsen at Hammel Neurorehabilitation Center for help with patient recruitment.

